# Testing of putative antiseizure drugs in a preclinical Dravet syndrome zebrafish model

**DOI:** 10.1101/2023.11.11.566723

**Authors:** P Whyte-Fagundes, A Vance, A Carroll, F Figueroa, C Manukyan, S.C. Baraban

**Author notes:** Corresponding author* Correspondence to Paige Whyte-Fagundes and Scott C. Baraban.

## Abstract

Dravet syndrome (DS) is a severe genetic epilepsy primarily caused by *de novo* mutations in a voltage-activated sodium channel gene (SCN1A). Patients face life-threatening seizures that are largely resistant to available anti-seizure medications (ASM). Preclinical DS animal models are a valuable tool to identify candidate ASMs for these patients. Among these, *scn1lab* mutant zebrafish exhibiting spontaneous seizure-like activity are particularly amenable to large-scale drug screening. Prior screening in a *scn1lab* mutant zebrafish line generated using N-ethyl-N-nitrosourea (ENU) identified valproate, stiripentol, and fenfluramine e.g., Federal Drug Administration (FDA) approved drugs with clinical application in the DS population. Successful phenotypic screening in *scn1lab* mutant zebrafish consists of two stages: (i) a locomotion-based assay measuring high-velocity convulsive swim behavior and (ii) an electrophysiology-based assay, using *in vivo* local field potential (LFP) recordings, to quantify electrographic seizure-like events. Using this strategy more than 3000 drug candidates have been screened in *scn1lab* zebrafish mutants. Here, we curated a list of nine additional anti-seizure drug candidates recently identified in preclinical models: 1-EBIO, AA43279, chlorzoxazone, donepezil, lisuride, mifepristone, pargyline, soticlestat and vorinostat. First-stage locomotion-based assays in *scn1lab* mutant zebrafish identified only 1-EBIO, chlorzoxazone and lisuride. However, second-stage LFP recording assays did not show significant suppression of spontaneous electrographic seizure activity for any of the nine anti-seizure drug candidates. Surprisingly, soticlestat induced frank electrographic seizure-like discharges in wild-type control zebrafish. Taken together, our results failed to replicate clear anti-seizure efficacy for these drug candidates highlighting a necessity for strict scientific standards in preclinical identification of ASMs.

## Introduction

Dravet syndrome (DS) is a devastating form of childhood epilepsy characterized by multiple clinical seizure types and increased early-life mortality. In these children, epilepsy presents in the first year of life with prolonged hemi-clonic or tonic-clonic seizures, followed by unprovoked seizures of varying etiologies including generalized tonic-clonic seizures ^1–3^. Increased risk of early mortality due to sudden unexpected death in epilepsy (SUDEP) is linked to DS. Quality-of-life (QOL) measures for DS patients (and caregivers) are an additional struggle as comorbidities include intellectual disability, developmental delays, movement and balance issues, language and speech deficits, sleep disturbance, and mood disorders. Currently available anti-seizure medications (ASMs) do not fully control seizures and polytherapy is common ^4^. Recent successes with stiripentol ^5^, fenfluramine ^6–9^, and cannabidiol (CBD) ^10^ provide hope - reducing seizures up to 60% in some patients - but each of these pharmacological treatments are associated with side effects or cardiovascular risk. Antisense oligonucleotides (ASOs) have shown exciting disease-modifying effect in DS mouse models and are currently in early clinical phase trials ^11^. However, despite these promising advances full seizure control and improved QOL remains an unmet medical need for most DS patients.

In over 80% of cases, DS is caused by *de novo* heterozygous *SCN1A* mutations resulting in haploinsufficiency. These include predicted loss-of-function (LOF) microdeletions, missense and truncation mutations ^12–14^. *SCN1A,* a gene encoding the voltage-gated sodium channel Na_v_1.1α subunits, is primarily expressed in parvalbumin-expressing fast-spiking inhibitory interneurons within the central nervous system ^15–17^. Reduced firing activity for a critical inhibitory interneuron sub-population providing feed-forward inhibition of excitatory neurons may contribute to the emergence of seizure activity in DS ^18^. While enhancement of Nav1.1 expression or activity on this interneuron sub-population may be an enticing therapeutic opportunity ^19–21^, it is also possible that this window for intervention is transient as studies in *Scn1a*^+/-^ mice have shown the impairment to resolve in early adulthood ^22^.

An important goal of preclinical research is to have well-characterized, reliable, and translatable animal models to facilitate discovery and development of novel DS treatments ^23,24^. Several existing preclinical DS mouse models have contributed to our understanding of the pathogenesis of this disorder and identification of potential therapeutics. Induced pluripotent stem cell (iPSC)-derived neurons from DS patients have further contributed to this literature ^25–27^, although unlike *in vivo* models these cell culture systems do not exhibit unprovoked seizure activity ^28^. In pursuit of this goal, we developed preclinical models based on zebrafish *scn1lab* mutation induced chemically ^23^ or using CRISPR/Cas9 gene editing ^29^. Homozygous *scn1lab* zebrafish mutants recapitulate haploinsufficiency (as a second zebrafish *scn1laa* gene is present) and replicate many clinically relevant phenotypes: (i) spontaneous unprovoked seizures, (ii) sleep disturbance, (iii) early fatality (iv) metabolic deficits and (v) pharmaco-resistance to three or more ASMs ^30^. Using *scn1lab* mutant zebrafish we screened more than 3000 drugs with a two-stage phenotype-based strategy incorporating sensitive behavioral and electrophysiological assays ^31^. Serotonin-acting drugs (fenfluramine, lorcaserin, clemizole and trazodone) as well as synthetic cannabinoids were identified in these preclinical screens ^23,31–33^. Both fenfluramine (Fintepla) and cannabadiol (CBD) were recently approved by the Federal Drug Administration (FDA) for treatment of DS validating this approach. Recently, additional drug candidates with putative antiseizure properties have also been reported. For example, intra-thalamic infusion of 1-EBIO reduced non-convulsive seizures in *Scn1a*^+/-^ mice ^34^ and pargyline reduced light-evoked brain activity in *scn1lab*^-/-^ zebrafish ^35^. However, as these discoveries were made in different laboratories, using different preclinical models and screening parameters, it is important to validate these discoveries in a scientifically rigorous manner before any translation to patients can be initiated.

## Materials and methods

### Zebrafish husbandry

All procedures described herein were performed in accordance with the Guide for the Care and Use of Animals (ebrary Inc., 2011) and adhered to guidelines approved by the University of California, San Francisco Institution Animal Care and Use Committee (IACUC approval: #AN171512-03A). Adult and juvenile zebrafish were maintained in a temperature-controlled facility on a 14-hour light and 10-hour dark cycle (9:00 AM to 11:00 PM PST). Juveniles (30 – 60 dpf) were fed twice daily, once with JBL powder (JBL NovoTom Artema) and the other with JBL powder mixed with live brine shrimp (Argent Aquaculture). Adults were also fed two times per day, first with flake food (tropical flakes, Tetramin) and then with flake food mixed with live brine shrimp). Zebrafish embryos and larvae were raised in an incubator kept at 28ᵒC on the same light/dark cycle as the facility in embryo medium consisting of 0.03% Instant Ocean (Aquariam Systems, Inc) in reverse osmosis-distilled water. Two *scn1lab* DS mutant zebrafish lines were utilized: ENU-generated *scn1lab*(*didy^s552^*) ^23,36^ and CRISPR-generated *scn1lab* (CRISPR mutant)^29^.

### Behavioral assay

Behavioral studies on 5 dpf larvae were conducted in a 96-well format with automated locomotion detection using a DanioVision system capturing at 25 frames per second and running EthoVision XT 11.5 software (DanioVision, Noldus Information Technology). *Scn1lab* mutant zebrafish larvae were sorted based upon pigmentation as mutants are much darker than wild types and, together with TLs, larvae were individually plated (1 per well) in 150 μL of embryo media and acclimated on the bench for 1 hour. Larvae were then placed in the DanioVision observation chamber and left undisturbed for a 20 minute habituation period. Baseline larval movements were tracked for 15 minutes, followed by a removal of 75 μL of media from each well and immediate addition of 75 μL of double concentrated candidate ASMs. Incubation with drugs occurred for 20 minutes prior to a tracking session for 15 minutes. Recording parameters included the following detection settings; subject: darker than background, method: differencing, sensitivity: ∼25-30, background changes: video pixel smoothing and medium slow, subject contour: 1 pixel with contour dilation set to erode first then dilate, subject size: minimum 15 and maximum 5178. For each trial, a minimum of 3 different clutches consisting of 10 larvae per condition were used. The max swimming velocity (mm/s) of each larva was calculated from baseline swimming conditions using a custom-written MATLAB script ^29^. Mean velocities (mm/s) presented for each larva are normalized to individual baseline swimming velocities.

### Survival and toxicity assay

Following each behavioral assay trial, 96-well plates of larvae were removed from the DanioVision and each larva was inspected under a Leica dissecting microscope for survival. To determine survival, larvae were monitored for intact circulation and visible heartbeat. Drug toxicity was established by initiating a touch response in each individual larva by gently contacting them with a pipette tip. If larvae failed to respond to touch, yet still had an intact circulation and heartbeat, the treatment was considered toxic, and the results were documented.

### Apoptosis assay

5 dpf WT and *scn1lab* larvae were incubated in acridine orange (AO) (10 μM, Sigma Aldrich, CAS: 65-61-2) solution for 20 minutes followed by three 5 minute washes in embryo media. During the entire protocol larvae were protected from light to maintain integrity of the AO. Following washes, larvae were mounted in 1% low melting point agar and pigmentation was removed from the tops of their heads to avoid obscuring the imaging area. Larvae were imaged under a Nikon C2 confocal at 10x magnification with a 488 laser for presence of apoptotic cells.

### Electrophysiology assay

Larvae were selected randomly from each behavioral/survival assay for follow up electrophysiological recordings. Two larvae at a time were cold anesthetized on ice in a custom recording chamber for 5 minutes and immobilized dorsal side up in 2% low melting point agarose (BP1260-100, Fisher Scientific). Recording chambers were placed on the stage of an upright microscope (Olympus BX-51W) and monitored continuously using a Zeiss Axiocam digital camera. Under visual guidance, gap-free LFP recordings (15 minute duration) were obtained from the optic tectum using a single-glass microelectrode (WPI glass #TW150 F-3) with approximately 1 μM tip diameter backfilled with 2 mM NaCl internal solution as described previously ^29,37–40^. LFP acquisition settings included low-pass filtering at 1 kHz, high-pass filtering at 0.2 Hz and sampling at 10 kHZ using a Digidata 1320 A/D interface (Molecular Devices). LFP data was stored on a PC computer running AxoScope 10.3 software (Molecular Devices) and scored for types of electrical activity as described in Griffin et al.^29^ Quantification of epileptiform events were performed blinded to the investigator with additional spectral analysis of seizure-like events utilizing MATLAB R2022b (MathWorks). Candidate ASMs were considered ‘positive hits’ if they were able to completely abolish all Type 2 activity in *scn1lab* mutant larvae without having any adverse side effects in TLs.

### Statistical analysis and reproducibility

Statistical tests were performed using GraphPad Prism. An unpaired t-test with Welch’s correction or One-way analysis variance with Dunnett’s multiple comparison test were used for determining significance of behavioral and toxicology assay data. Kruskal-Wallis with Dunn’s multiple comparisons test or One-way analysis variance with Dunnett’s multiple comparison were used for electrophysiological data. Any data presented in heatmap form are presented as a mean, otherwise all individual values are presented along with the distribution of data or the mean ± S.E.M. All data was collected and analysis was performed blinded to the genotype.

### Pharmacology

All candidate ASMs were dissolved in 100% DMSO (Dimethyl sulfoxide, Sigma Aldrich, CAS: 67-68-5) to a final stock concentration of 100 mM. Each ASM was diluted in embryo media to double concentrated working solutions (i.e. 500 μM, 200 μM and 20 μM) for final assay concentrations resulting in 250 μM with 0.25% DMSO (high concentration), 100 μM with 0.1% DMSO (medium concentration) and 10 μM with 0.01% DMSO (low concentration). Note, no differences in larval phenotypes are seen with DMSO at concentrations below 1% ^41^. Candidate ASM order information was as follows: 1-Ethyl-2-benzimidazolinone (Alternative name 1-EBIO, Tocris, CAS: 1041), AA43279 (Focus biomolecules, CAS: 354812-16-1), Chlorzoxazone (Sigma Aldrich, CAS: 95-25-0), Donepezil hydrochloride (Sigma Aldrich, CAS: 120011-70-3), Lisuride maleate (Tocris, CAS: 40-5210), Mifepristone (Sigma Aldrich, CAS: 84371-65-3), Pargyline hydrochloride (Sigma Aldrich, CAS: 306-07-0), SAHA (Alternative name Vorinostat, Sigma Aldrich, CAS: 149647-78-9), Soticlestat (DC Chemicals, CAS: 1429505-03-2). It is important to note that Mifepristone, an antiprogestogen identified in Eimon et al.^35^ precipitated out of solution and was not included in the screening assays.

## Results

We used *scnl1lab* loss-of-function CRISPR-Cas9-generated zebrafish carrying a 7 base-pair deletion ^29^ and ENU-generated zebrafish carrying a T-to-G mutation ^36^. Seizure frequency monitored using single-electrode local field potential (LFP) recording was comparable for CRISPR- and ENU-generated *scn1lab* zebrafish lines (**Supplementary Fig. 1a**). Using acridine orange (AO) staining for apoptosis ^42^, *scn1lab* mutant zebrafish do not show increased levels of AO staining compared to wild-type (WT) sibling controls (**Supplementary Fig. 1b**). Herein, we used both *scn1lab* mutant zebrafish lines to test drug candidates recently identified as exhibiting putative anti-seizure activity in preclinical DS models (**Table I**).

**Table 1:** Candidate drugs identified in preclinical DS models. Proposed drugs in this table are purported to have anti-seizure properties. Mechanisms of action for each candidate are indicated.

### Locomotion-based screening of putative antiseizure drugs in scn1lab zebrafish

Single-point locomotion tracking is a rapid surrogate method to monitor behavioral seizure activity in freely swimming zebrafish larvae. On average, *scn1lab* larvae reach significantly higher maximum swimming velocities [above 25 mm/sec^29^] compared to WTs. Plotting individual larval velocity over time, these swim activities appear in burst-like patterns followed by brief periods of lower velocities (**Fig. 1a**). Here, *scn1lab* mutant and WT sibling control larvae were tracked in a 96-well plate format. Candidate drugs were added to each individual well at three different concentrations: 10, 100 or 250 μm (**Fig. 1b**). AA43279, a Nav1.1 sodium channel activator ^43^, increased swim velocities in WT larvae at 100 μM; AA43279 did not alter swim velocities for *scn1lab* mutants at any concentration tested. Donepezil, a reversible acetylcholinesterase inhibitor ^44^, increased swim velocities for WT larvae at 10 and 100 μM and for *scn1lab* mutant larvae at 100 and 250 μM. 1-EBIO (a calcium-activated potassium channel activator; ^45^) at 100 and 250 μM, chlorzoxazone (a muscle relaxant; ^46^) at 10, 100 and 250 μM, and lisuride (a serotonin receptor agonist; ^47^) at 10, 100 and 250 μM, significantly reduced swim velocities in *scn1lab* mutants. However, reductions in swim velocities were also seen for all three drugs in WT control larvae. (**Fig. 1b**). Representative tracking plots are shown for two larvae treated with each drug candidate that either significantly increased (**red**) or decreased (**blue**) average swim velocity of WT control (**Fig. 1c**) or *scn1lab* mutant (**Fig. 1d**) larvae, respectively. For each of these drugs, normalized velocities of each treated larvae are plotted alongside vehicle (0.25% DMSO) treated control larvae in **Figures 1c** **& 1d**.

**Figure 1:**
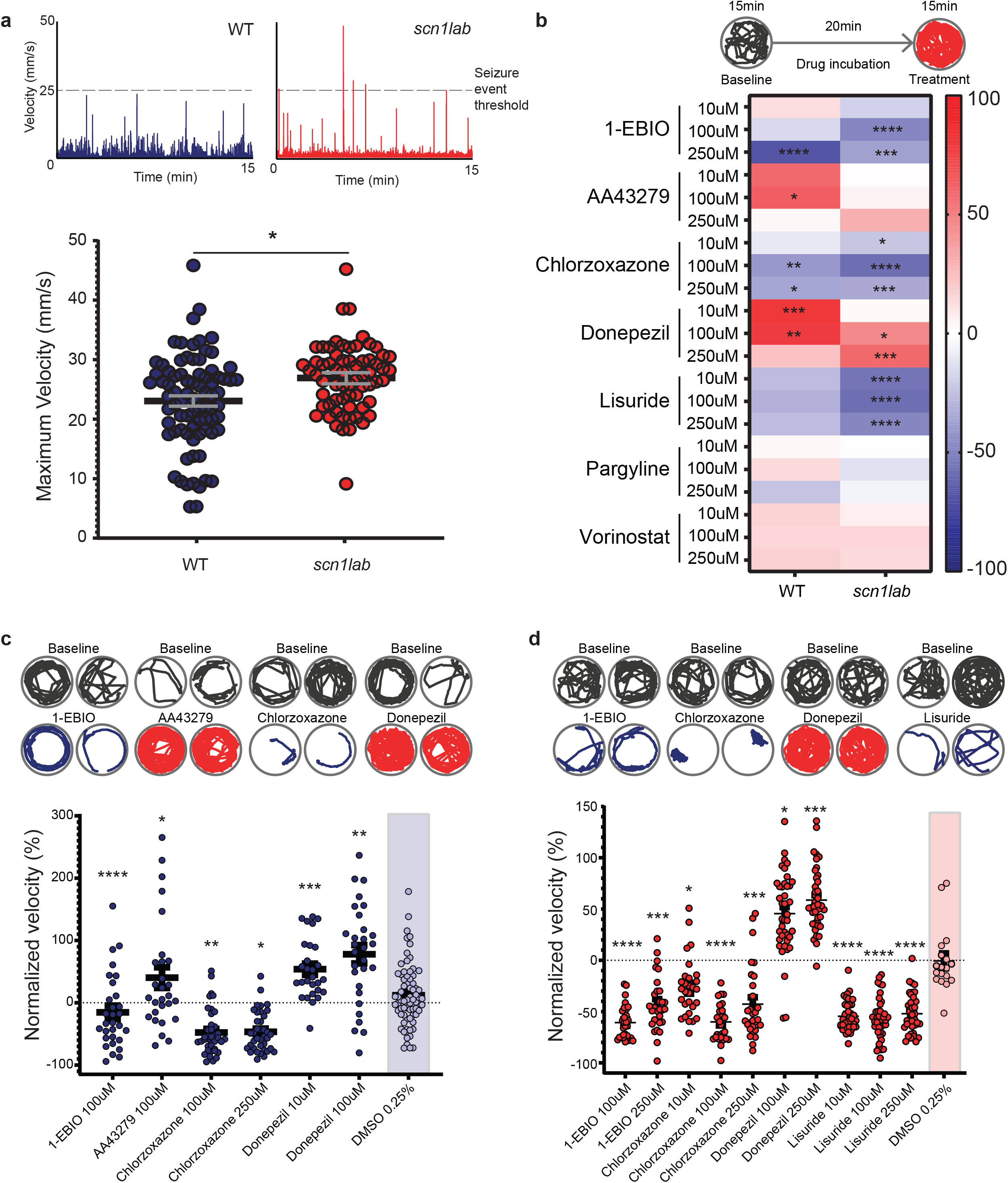
Behavioral phenotypic screening of candidate ASMs in WT and scn1lab larvae. **(A)** Representative baseline swim velocities (mm/s) over time for 5 dpf WT (blue) and scn1lab (red) mutant larvae. Threshold for behavioral seizure events indicated by dotted line (28 mm/s). Scn1lab larvae (N = 60) have significantly (p = 0.014) higher maximum velocities on average compared to WTs (N = 70). **(B)** Timeline for behavioral tracking acquisition (top). Heatmap of the percent change in average swimming velocity from baseline after treatment with candidate ASMs at three different concentrations. Significant changes from vehicle control are indicated by stars with an N = 30 or 40 per condition across minimum 3 trials for WT and scn1lab larvae. **(C)** Behavioral tracking plots for WT larvae showing baseline activity followed by ASM treatment for drugs that caused a significant change in swim velocity compared to DMSO controls. The plot below highlights the normalized velocity in percent, after ASM treatment for each larvae recorded. AA43279 significantly increased swim velocity in WT larvae at 100 µM (N = 40, p < 0.0001). **(D)** Behavioral tracking plots for scn1lab larvae showing baseline activity followed by ASM treatment for drugs that caused a significant change in swim velocity compared to DMSO controls. Donepezil increased swim velocities for scn1lab larvae (N = 40, p < 0.01). 1-EBIO, chlorzoxazone and lisuride significantly reduced swim velocities (N = 30 – 40, p < 0.05-0.0001). The plot below highlights the normalized velocity in percent, after ASM treatment for each larvae recorded. p<0.05 = *, p<0.005=**, p<0.0005=***, p<0.0001=****.

To assess potential toxic or sedative actions, following each behavioral assay all larvae were monitored under a dissecting microscope to determine presence of a heartbeat (indicative of survival) and probed to establish responsiveness to touch ^48^. Larvae mostly survived treatment with each drug candidate regardless of genotype. However, 100 μM vorinostat (an HDAC inhibitor ^49^) significantly reduced survival in *scn1lab* larvae (**Fig. 2a**). WT control larvae were uniquely non-responsive to pargyline and donepezil at 100 μM (**Fig. 2b**); *scn1lab* mutants lacked touch responses following vorinostat treatment at 100 μM (**Fig. 2c**). 1-EBIO and chlorzoxazone significantly reduced larval responses in WT control and *scn1lab* mutant larvae (**Fig. 2d**) ^49^.

**Figure 2:**
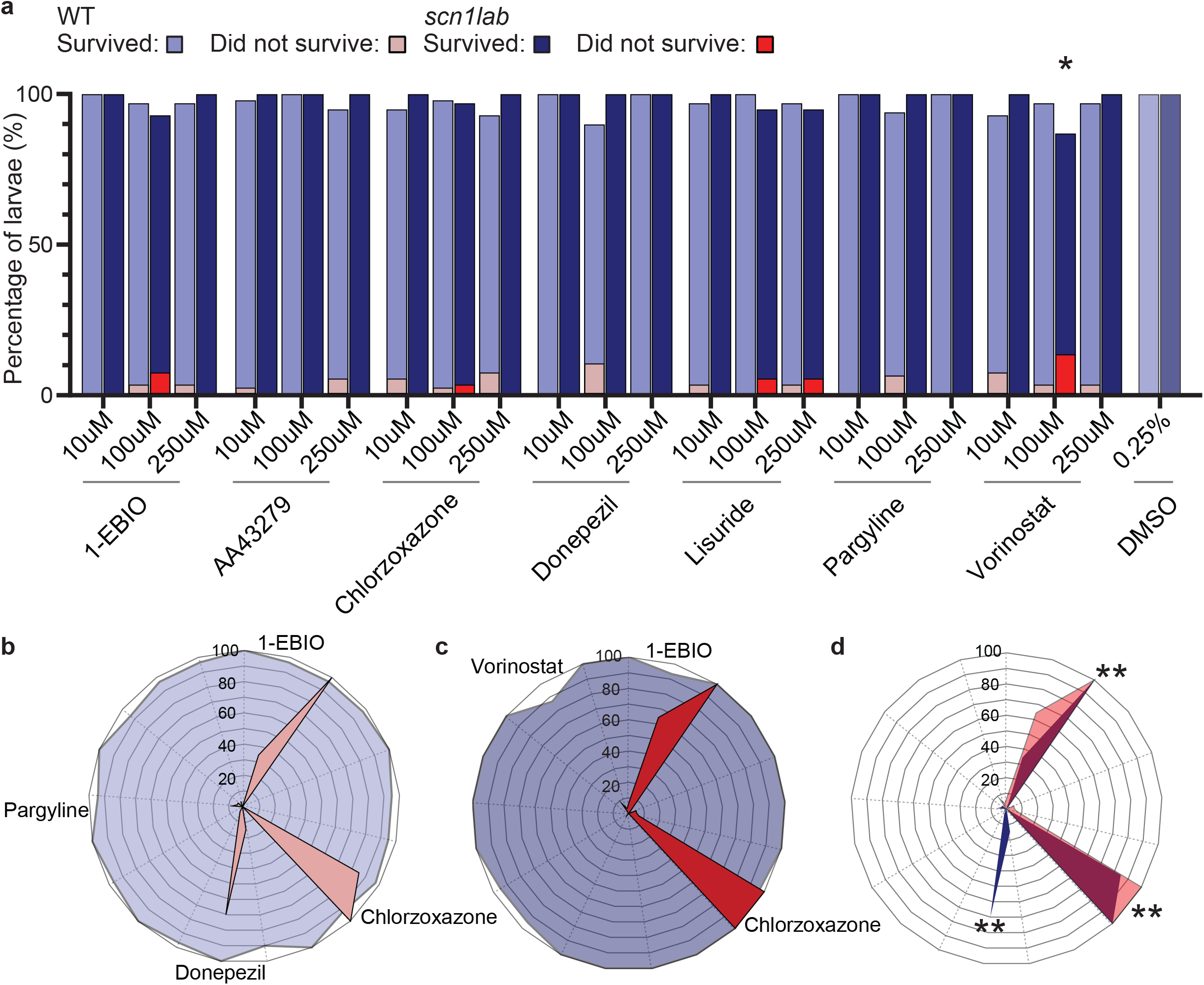
Toxicology assay for larvae following behavioral assessment. **(A)** Percent survival of WT (light blue) and scn1lab (dark blue) larvae (N = 30 or 40 per condition with minimum independent 3 replicates). Percentage of larvae that did not survive are overlayed, WT in light red and scn1lab in dark red. Vorinostat at 100 µM significantly reduced scn1lab survival compared to DMSO treated controls. **(B)** Radar plot quantifying touch responses of WT larvae. Dotted lines segment each candidate ASM and toxicity is plotted as a percentage of larvae. Toxic ASMs are labelled. **(C)** Radar plot showing candidate ASM toxicity for scn1lab larvae. **(D)** Radar plot of WT and scn1lab toxicity overlayed. 1-EBIO and Chlorzoxazone were toxic for both populations. Donepezil uniquely impacted WT larvae. One-way ANOVA was performed for statistical analysis, p<0.01 = *, p < 0.0001 = **.

### Electrophysiology-based screening of putative antiseizure drugs in scn1lab zebrafish

Representative *in vivo* LFP recordings for type 0, type 1 and type 2 electrical activities from untreated larvae are shown in **Figure 3a**; traces scored as type 0 are considered normal baseline electrical activity. Type 1 events are considered interictal-like activities and type 2 are considered ictal- or seizure-like electrographic events, which manifest as large voltage deflections with poly-spiking typically followed by a post-ictal depression. As expected, WT control larvae recorded during basal conditions revealed only type 0 events (**Fig. 3a**). The plot showing distribution of electrical events in *scn1lab* larvae confirms that the majority of mutant larvae exhibit at least one spontaneous type 2 event within a 15 min LFP recording epoch (**Fig. 3a**).

**Figure 3:**
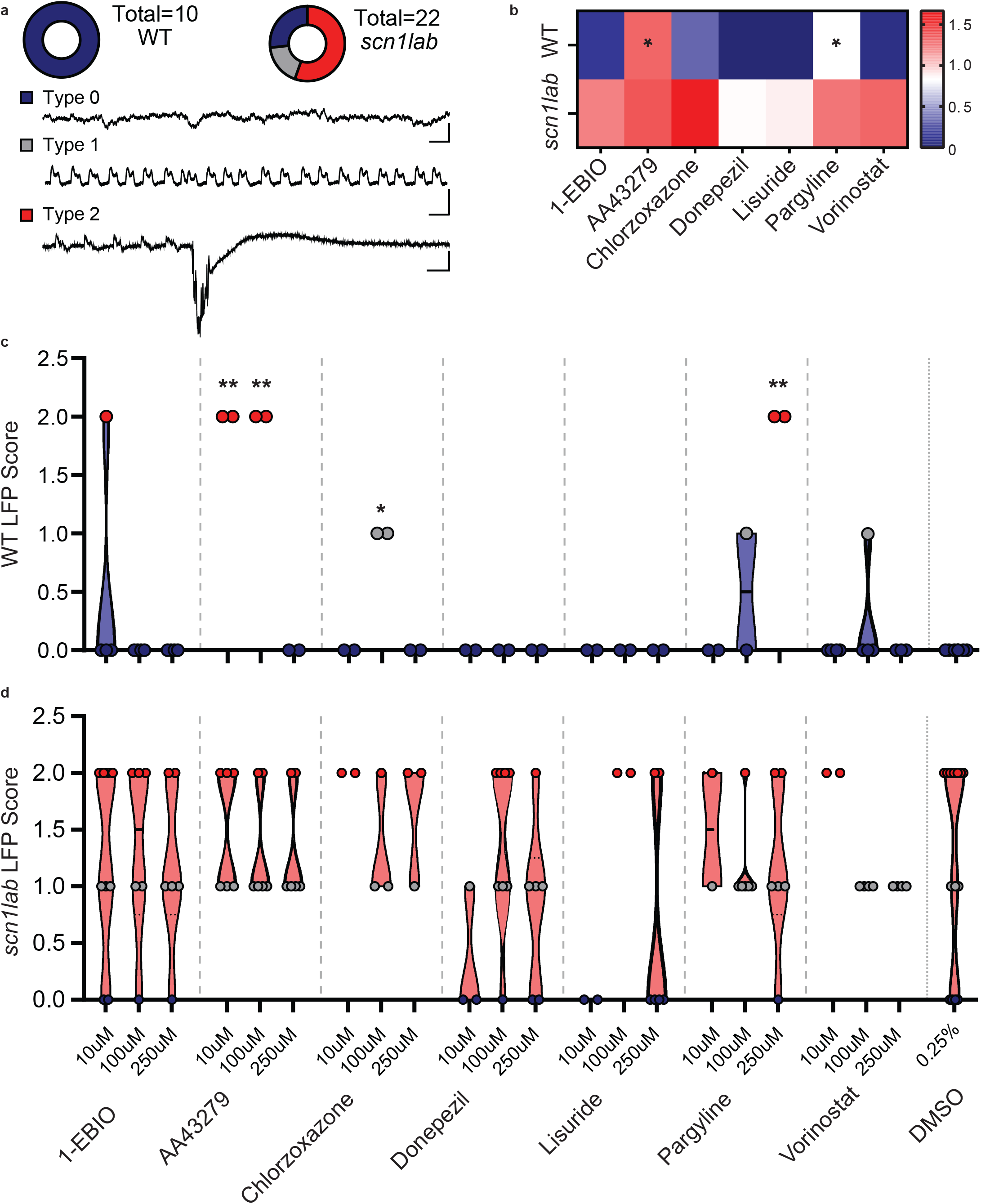
Electrophysiological recordings of larvae treated with candidate ASMs. **(A)** Donut plots of untreated WT (left) and scn1lab (right) larvae showing the distribution of type 0 (normal), type 1 (interictal-like) and type 2 (ictal-like) electrical activity along with representative LFP traces. **(B)** Heatmap of average LFP scores for both WT and scn1lab larvae after treatment with candidate ASMs at 3 different concentrations showing a significant increase in activity for WT larvae treated with AA43279 (N=7, p =0.01) and pargyline (N = 7, p = 0.05). **(C)** Violin plot showing scored electrophysiological recordings from individual WT larvae aftertreatment with candidate ASMs at each concentration. AA43279, Chlorzoxazone, and Pargyline significantly induced abnormal activity in WT larvae compared to control DMSO treatment (N = 2-4 per condition, p<0.0001). 1-EBIO induced type 2 and vorinostat induced type 1 events in 1 out of 4 larvae. **(D)** Violin plot showing scored electrophysiological recordings from individual scn1lab larvae. No drugs reliably prevented seizure activity compared to control DMSO treated larvae, with only some drugs having a mild modulatory effect on seizure activity (N = 2-6 per condition). Statistical tests include Kruskal-Wallis and One-way ANOVA, p<0.05 = *, p < 0.0001 = **, all other data was not significantly different. Scale for traces are 1s by 0.1mV.

To first gain an understanding of the overall effect candidate ASMs had on larvae, all electrophysiology traces were blindly scored as type 0, 1 or 2. Next, these qualitative scores were averaged across pooled recorded larvae for each ASM drug candidate. In WT controls, the majority of candidate ASMs did not alter LFPs compared to DMSO-treated controls, except for AA43279 and pargyline which showed signs of significantly increased electrical activity (**Fig. 3b, top**). In *scn1lab* mutants, no significant evidence of reduced electrical activity was noted for any of the nine drug candidates. Although donepezil and lisuride did appear to modulate occurrence of some type 2 events this did not reach a statistically significant level (**Fig. 3b, bottom**). LFP scores for each recorded WT larvae were then plotted to determine which concentrations of AA43279 and pargyline were altering electrical activity (**Fig. 3c**). Surprisingly, AA43279 (10 and 100 μM), and pargyline (250 μM) evoked type 2 ictal-like events in WT controls. Although 1-EBIO, chlorzoxazone and lisuride were identified as ‘positive hits’ in the first-stage behavioral assays, they failed to abolish electrographic type 2 ictal-like events in LFP recordings from *scn1lab* zebrafish mutants (**Fig. 3d**).

### Paradoxical effects of soticlestat in zebrafish

Soticlestat, a cholesterol 24-hydroxlase inhibitor, elevated thresholds for hyperthermia-induced tonic-clonic seizures in two haploinsufficient *Scn1a* mouse models ^50^ and is currently an investigational candidate as an adjunctive therapy for DS ^51^. To further evaluate this candidate, we first examined spontaneous swim behavior in WT and *scn1lab* mutant larvae exposed to increasing concentrations of soticlestat. Surprisingly, soticlestat increased swim velocities for WT larvae at concentrations of 100 and 250 μM (**Figs. 4a-b**). Soticlestat also decreased survival rates of WT larvae with increasing drug concentration with no clear impact on *scn1lab* mutant larvae (**Fig. 4b**). Next, we performed electrophysiology recordings. Representative electrographic seizure activity and a corresponding spectrogram are shown in Figure 4c. Soticlestat consistently induced type 1 interictal-like and 2 ictal-like activity in WT larvae at all concentrations tested (**Fig. 4f**) but failed to abolish electrographic seizure activity in *scn1lab* mutant larvae (**Fig. 4g**).

**Figure 4:**
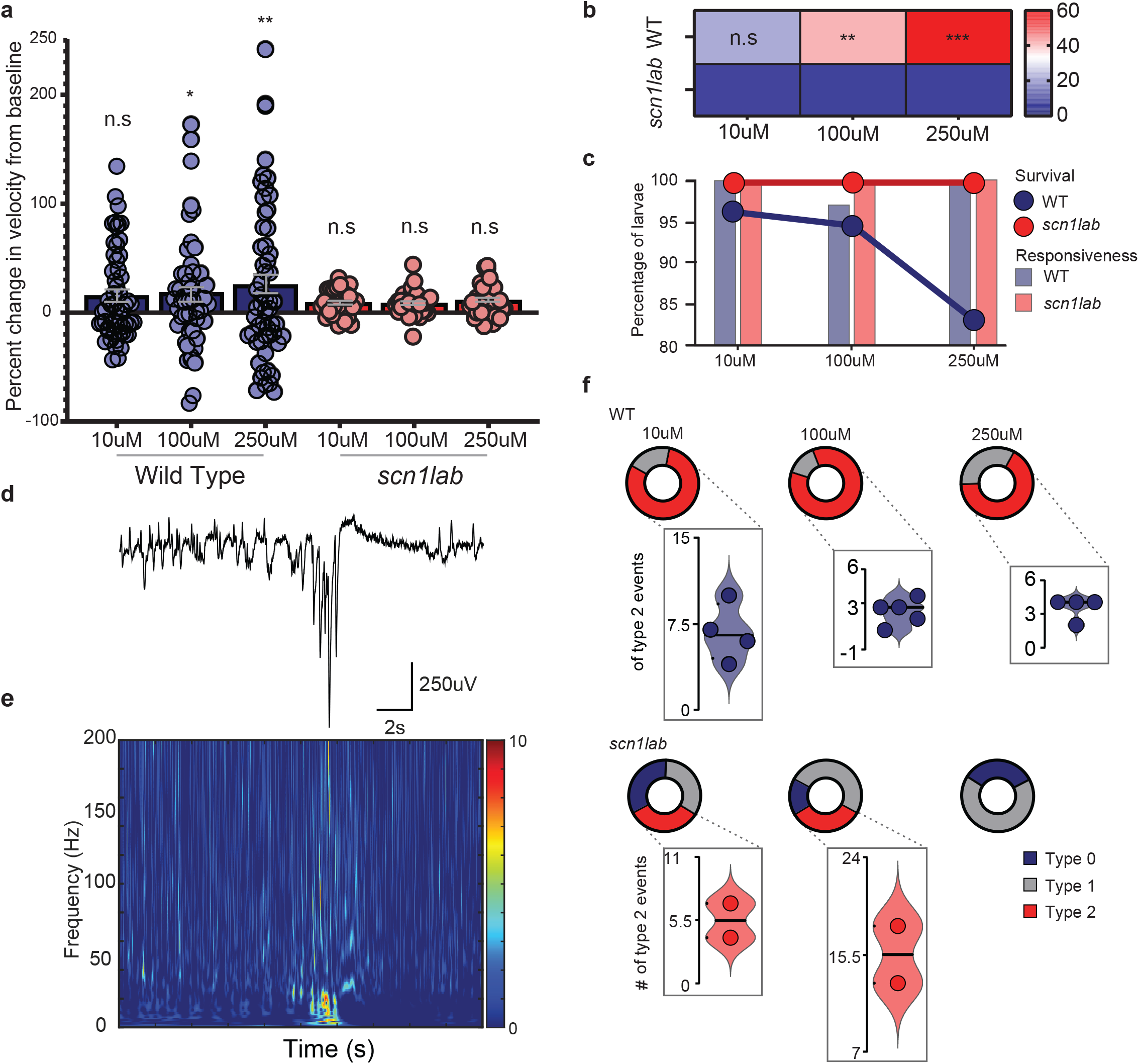
In depth assessment of soticlestat treatment of seizure activity for WT and scn1lab larvae. **(A)** Swimming behavior for individual WT (Blue, N = 60) and scn1lab (Red, N = 40) larvae after treatment with 3 concentrations of soticlestat represented as a percent change in velocity from baseline. Increasing concentrations of soticlestat significantly increased velocity of WTs compared to DMSO treated controls but did not alter the velocity of scn1lab larvae at any concentration. **(B)** Heatmap showing significant increases in average normalized swimming velocities of soticlestat treated WT larvae compared to scn1lab larvae. **(C)** Increasing concentrations of soticlestat treatment decreased survival rates of WT larvae but did not impact scn1lab larval survival or responsiveness. **(D)** LFP recording sample of seizure activity induced in WT larvae after soticlestat treatment along with **(E)** associated spectrogram. Note the high-frequency activity and electrodecremental LFP response following an ictal-like event. **(F)** Donut plots representing the percentage of fish showing type 0, 1 and 2 activity for both WT (top) and scn1lab (bottom) larvae treated with 3 concentrations of soticlestat with quantification of type 2 events for each seizing fish below each plot. Soticlestat induced type 1 and 2 activity in WT larvae at all concentrations but did not abolish seizure activity in scn1lab larvae unless at high concentrations. Unpaired t-test was performed for statistical analysis, p<0.05 = *, p<0.01= **, p < 0.0001 = **.

## Discussion

Preclinical SCN1A experimental animals, primarily in mice and zebrafish, provide insights into the pathogenesis and potential treatment of DS. While a *scn1lab* zebrafish line successfully identified ‘standard-of-care’ ASMs used in DS patients (e.g., fenfluramine, stiripentol and valproic acid), as well as several serotonin receptor agonists with compassionate-use efficacy in this patient population (e.g., lorcaserin and trazodone), *Scn1* mice have failed to identify these drugs ^28^. As preclinical DS drug discovery programs vary in the types of animals (zebrafish vs. mice) or protocols (hyperthermia-induced, light-induced or spontaneous seizures) it has become imperative to set strict standards for accurately identifying and validating preclinical drug discoveries. Here, we used our validated two-stage screening platform to evaluate nine anti-seizure drug candidates described in recent preclinical mouse (l-EBIO, chlorazoxazone and donepezil) or zebrafish (lisuride, AA43279, vorinostat, and pargyline) models. Using phenotypic behavioral and electrophysiological zebrafish assays we were unable to replicate anti-seizure activity reported for any of these drugs. Surprisingly, we also discovered that soticlestat, currently in Phase 3 clinical trials (clinicaltrials.gov), evoked electrographic seizure activity in otherwise healthy wild-type zebrafish.

Cannabadiol and fenfluramine, recently approved as adjunctive ASMs for DS, significantly reduce seizure frequency in patients ^9,52^. Preclinical identification of these FDA-approved drugs in Scn1 mouse models has been difficult. Using hyperthermia-induced seizure protocols, cannabadiol attenuated seizure severity in *Scn1a^+/-^* mice at concentrations above 100 mg/kg ^53^ or decreased seizures in combination with clonazepam ^54^ but failed to show anti-seizure activity in *Scn1a^A1783/WT^* mice ^55^. Fenfluramine, at concentrations as high as 25 mg/kg, failed to show protection against hyperthermia-induced seizures in *Scn1a^A1783/WT^*mice ^55^. In contrast, five different synthetic cannabinoids [note: cannabadiol is oil-based and short-term exposure is toxic to zebrafish larvae ^56^] reduced high-velocity seizure-like behaviors and suppressed electrographic seizure activity in *scn1lab* (*didy^s552^*) zebrafish mutants ^31^. In addition to highlighting the predictive pharmacological validity of *scn1lab* zebrafish models these studies also emphasize the importance of incorporating multiple outcome measures e.g., toxicity, behavior and electrophysiology. Here, in locomotion assays using both WT and *scn1lab* mutant zebrafish, we noted a reduction in swim activity with lisuride, chlorzoxazone and 1-EBIO. Based on subsequent toxicology assessments of survival and touch responses, we attribute ‘false positive’ behavioral hits on SK channel activators 1-EBIO and chlorzoxazone to toxicity. Meanwhile, lisuride, which reduced locomotor activity in *scn1lab* mutant zebrafish, may be attributed to a sedative or toxic action ^57–59^. On the other hand, donepezil dramatically increased swim velocity in WT zebrafish. WT zebrafish larvae are routinely employed in high-throughput behavioral screens for evaluating drug toxicity ^60–62^. Cardiac morphology or automated functional imaging assays using WT zebrafish larvae offer another layer of toxicology screening ^63,64^. In our small shelf screen, we added a sensitive electrophysiology assay to evaluate potential CNS side-effects. Interestingly, LFP recordings from WT zebrafish larvae exposed to soticlestat suggest an alteration of brain activity that conservatively indicates hyperexcitability and more cautiously suggests a level of seizure-like activity.

Although DS patients are often treated with polypharmacy, cannabinoids and serotonin signaling pathways have recently emerged as potential mechanism(s) for seizure control. Here, we tested putative anti-seizure candidates with a wide range of mechanisms from cholesterol 24-hydroxylase, monoamine oxidase, glucocorticoid hormone receptor or histone deacetylase inhibitors to ion channel modulators (see **Table I**). Assays and models used to discover these drug candidates were also varied. In two different *scn1lab* mutant zebrafish lines, Eimon et al.^35^ employed a high-throughput light-stimulus assay and custom-made electrophysiology platform to identify pargyline and mifepristone. Whether light-evoked activity represents epilepsy is unclear, and although both *scn1lab* zebrafish lines exhibit spontaneous seizures ^23,29^ these drugs were not tested against this type of epileptic activity. Weuring et al.^65^ identified the Nav1.1 channel activator AA43279 using a CRISPR-generated *scn1lab* zebrafish line exhibiting spontaneous short, small-amplitude spike events recorded using an electrode placed externally on top of the forebrain. AA43279 reduced the frequency of these electrical events by approximately 50% but was less effective when compared to fenfluramine. A prolonged 22 hour exposure to lisuride (an anti-Parkinson drug with both dopamine and serotonin binding activity) effectively abolished locomotor activity in *scn1lab* (*didy^s552^*) mutants and reduced epileptiform activity in electrophysiology recordings by approximately 33%. However, toxicology and electrophysiological assessments in the data presented here highlight that prolonged drug exposures may confound screening result outcomes (also see ^28^). Vorinostat (Zolinza), an FDA-approved HDAC inhibitor, initially discovered using larval zebrafish bioenergetics assays suppressed interictal-like (type 1) events in a *kcna1*-morphant zebrafish and was briefly discussed as a potential clinical trial candidate for DS but only slightly modified *scn1lab* larval electrical activities at high concentrations ^66^. In *Scn1a^+/-^* mice, donepezil was shown to be protective against hyperthermia-induced seizure and 1-EBIO reduced non-convulsive EEG seizure activity ^67^. Based on the latter findings, Ritter-Makinson et al.^34^ proposed that another FDA-approved SK-channel activator (chlorazoxone) could be a novel therapy candidate for DS patients. It is, perhaps, not entirely surprising that drug candidates discovered in mouse models may not be the best choice for translation to humans, as it is estimated that up to 95% of drugs identified in mice ultimately fail in clinical trials ^68–70^.

Although zebrafish are often over-looked as a vertebrate model for ASM discovery, there is emerging interest in finding new drug candidates for rare epilepsies using zebrafish. Here we confirm and extend our previously published protocols using single-point locomotion tracking and LFP recordings in a *scn1a* zebrafish model to evaluate a series of recently discovered drug candidates. Unfortunately, we were unable to provide clear evidence of anti-seizure activity for any of these drugs. Moreover, careful testing in wild-type larvae uncovered potential issues with donepezil, 1-EBIO, chlorazoxone and soticlestat that warrant further study. Taken together, the present study establishes a foundation for rigorous preclinical screening of drugs using *scn1lab* zebrafish.

## Data availability

Data sets generated during the current studies are available from corresponding authors upon reasonable request.

## Contributions

P.W.F and S.C.B conceived the project. P.W.F, and A.V collected behavioral and toxicity data.

F.F and C.M performed preliminary screening and replicates on the previous version of the Danio Vision. P.W.F, A.V and A.C performed apoptosis assays. P.W.F analyzed the behavioral and toxicology data. P.W.F collected and analyzed electrophysiology data. P.W.F and S.C.B wrote and edited the manuscript.

