## Supplementary figures and images for "Testing of putative antiseizure drugs in a preclinical Dravet syndrome zebrafish model"

### Supplementary figure 1

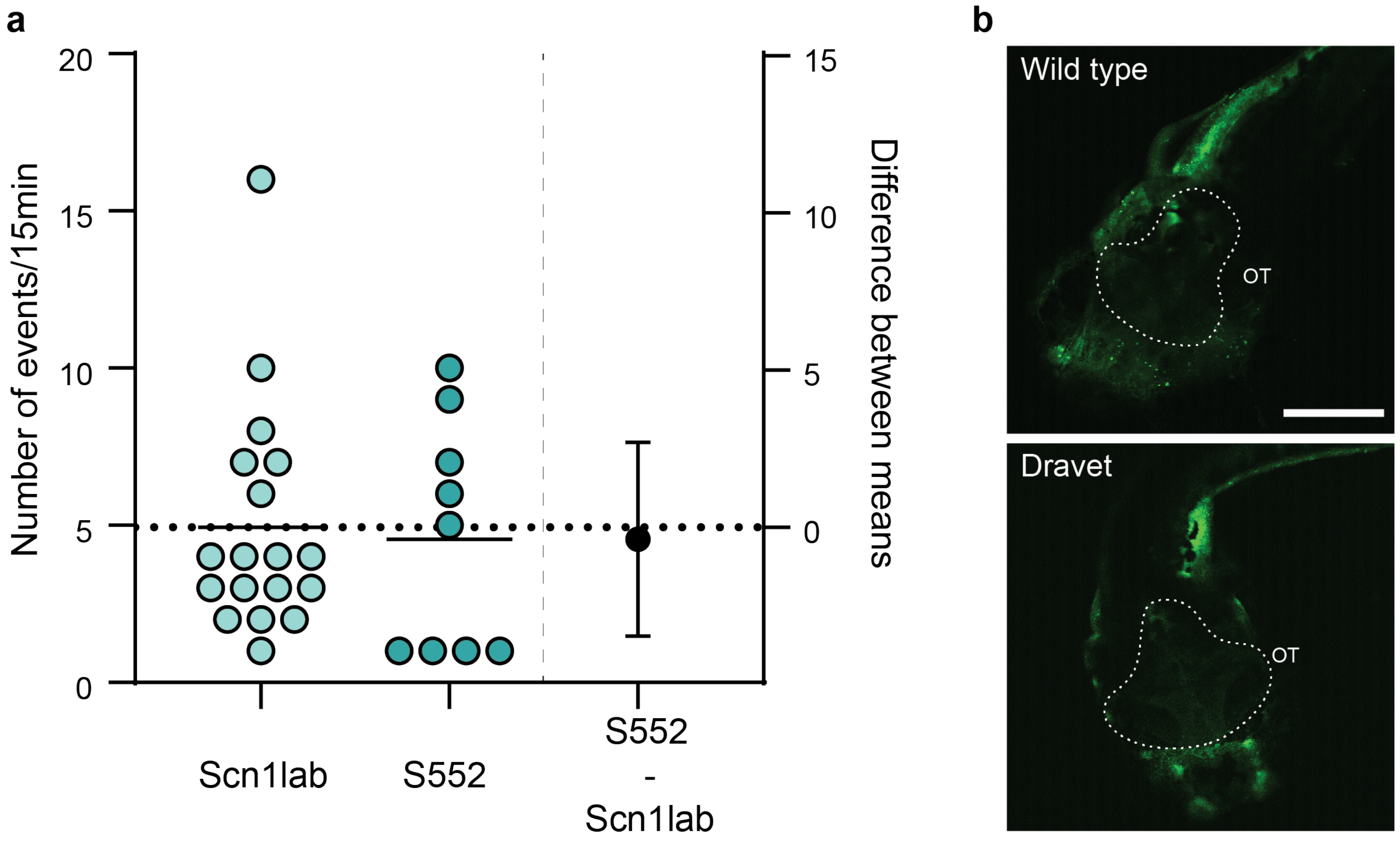
