## Supplementary table 1 for "Testing of putative antiseizure drugs in a preclinical Dravet syndrome zebrafish model"

**Table 1: Candidate drugs identified in preclinical DS models.** Proposed drugs in this table are purported to have anti-seizure properties. Mechanisms of action for each candidate are indicated.

| **Candidate drugs** | **Proposed mechanisms** |
|---|---|
| 1-EBIO | Activator of calcium activated potassium channels |
| AA43279 | Sodium (Nav1.1) channel activation |
| Chlorzoxazone | Activator of calcium activated potassium channels |
| Donepezil | Cholinesterase inhibitor |
| Lisuride | Serotonin receptor agonist |
| Mifepristone | Glucocorticoid hormone inhibitor |
| Pargyline | Selective monoamine oxidase-B inhibitor |
| Soticlestat | Cholesterol 24-hydroxylase inhibitor |
| Vorinostat | Broad histone deacetylase inhibitor |

*1-EBIO: 1-ethyl-1,3-dihydro-2H-benzimidazol-2-one
